# GPR15 in colon cancer development and anti-tumor immune responses

**DOI:** 10.1101/2021.03.13.435189

**Authors:** Hong NamKoong, Bomi Lee, Gayathri Swaminathan, Seong-Joon Koh, Stephan Rogalla, David Mikhail, Aida Habtezion

**Author notes:** Corresponding Author: Aida Habtezion, MD, MSc, Alway Building, Room M211, 300 Pasteur Drive, Stanford, CA 94305. Equal contribution (Hong NamKoong, Bomi Lee).

## Abstract

G protein-coupled receptor 15 (GPR15) is a chemoattractant receptor which in response to its ligand, C10orf99/GPR15L, promotes colon homing of T cells in health and colitis. The functional role of GPR15 in colon cancer is largely unexplored, motivating our current studies using murine colon cancer models and human colorectal cancer (CRC) tissues. Our initial analysis of human CRC specimen revealed significant reduction in GPR15 expression and frequency of GPR15^+^ immune cells in tumors compared to ‘tumor-free’ surgical margins. In the AOM/DSS murine model of colitis associated colon cancer (CAC), we observed increased colonic polyps/tumor burden and lower survival rate in *Gpr15*-deficient (KO) compared to *Gpr15*-sufficient (Het) mice. Analysis of immune cell infiltrates in the colonic polyps showed significantly decreased CD8^+^ T cells and increased IL-17^+^ CD4^+^ and IL-17^+^ CD8^+^ T cells in *Gpr15*-KO than in Het mice. GPR15 deficiency thus alters the immune environment in colonic polyps to mitigate T cell-mediated anti-tumor responses resulting in severe disease. Consistent with a protective role of GPR15, administration of GPR15L to established tumors in the MC38 CRC mouse model increased CD45^+^ cell infiltration, enhanced TNF*α* expression on CD4^+^ and CD8^+^ T cells at the tumor site and dramatically reduced tumor burden. Our findings highlight an important, unidentified role of the GPR15-GPR15L axis in promoting a tumor-suppressive immune microenvironment and unveils a novel, colon-specific therapeutic target for CRC.

## Introduction

CRC is a complex, heterogenous disease and is a leading cause of cancer-related deaths world-wide^1^. CRC is associated with a multitude of risk factors such as obesity, low-fiber and high-fat diet, heavy alcohol consumption, smoking, family history, inflammatory bowel disease (IBD) and so on^2–4^. Patients diagnosed with chronic colitis, such as Crohn’s disease (CD) and ulcerative colitis (UC), are prone to developing a distinct type of CRC, termed colitis associated colon cancer (CAC)^5–7^. Chronic, relapsing inflammation results in severe intestinal damage and sets the stage for colon cancer development by shaping a pro-tumor immune microenvironment. The intricate balance between tumor infiltrating immune cells including effector cells such as T cells, NK cells, immune-suppressive regulatory T cells (Tregs), myeloid cells, dendritic cells, B cells and their interactions with cellular and molecular components in the tumor microenvironment (TME) influence tumor development and clinical outcomes. For instance, a prominent Th1/Th2 phenotype shift, expansion of myeloid-derived suppressor cells (MDSC), tumor-associated macrophages (TAMs), decreased CD8^+^ T cells along with increased representation of certain inflammatory mediators such as IL-6, IL-17 contribute to tumor advancement. Conversely, infiltration of tumor-fighting cytotoxic CD8^+^ T cells, Th1 cells and generation of mediators such as IFN*γ*in the TME have been shown to collaborate in mounting anti-tumor responses resulting in cancer regression^8^. Several studies highlight a positive correlation between increased infiltration and high densities of CD3^+^ T cells (effector CD4^+^ or CD8^+^ T cells) at tumor sites to disease-free survival in CRC patients^9–12^. Interaction between chemokine and chemokine receptors facilitate traffic of T cells to the tumor. Pre-clinical development of modulators of chemokines and chemokine receptors is being actively pursued to be used as stand-alone or combination therapy for various oncology indications including CRC^13^.

GPR15 (also known as BOB) is seven-transmembrane domain, class A G-protein coupled receptor (GPCR), first identified as a co-receptor for human immunodeficiency virus (HIV) or simian immunodeficiency virus (SIV)^14–16^. GPR15 is expressed on regulatory and effector CD4^+^ T cells, fetal thymic epidermal T cells, B cells and plasmablasts. Studies from our group and others have identified GPR15 as a mucosal chemoattractant/trafficking receptor that facilitates migration of effector and regulatory T cells to sites of inflammation in the large intestine and plays a crucial role in colitis pathogenesis^17^. GPR15 was deorphanized with the identification of its cognate chemoattractant ligand, named GPR15L, encoded by the human gene *C10orf99* and mouse *2610528A11Rik*. GPR15L is expressed in human and mouse epithelial cells exposed to the environment, in organs such as the colon and skin^18, 19^. In contrast to the well-studied role of GPR15 in colitis and other inflammation conditions, the function of GPR15-GPR15L signaling axis in CRC is poorly understood. In particular, experimental evidence linking immune functions of GPR15 and GPR15-mediated mechanisms to CRC development is lacking.

In this report, we delineate a novel tumor-suppressive function of GPR15 using CRC mouse models and patient samples. We demonstrate using human colon cancer samples that GPR15 expression and GPR15^+^ T cells are significantly reduced in tumors compared to surgical tumor margins (STM), which are regarded ‘tumor-free’. In an AOM-DSS mouse model of CAC, *GPR15* deficiency resulted in increased tumor burden, morbidity and mortality concomitant with reduced T cell infiltration, especially CD8^+^ T cells in the tumor microenvironment. The immune infiltrates in the colonic polyps of *Gpr15-*KO mice as well as human CRC tumor samples with reduced GPR15 expression showed significant increases in the population of pro-inflammation mediators in the tissue microenvironment such as IL-17^+^ CD4^+^ and IL-17^+^ CD8^+^ T cells Consistent with a protective role of the GPR15, administration of GPR15L to established tumors in the MC38-CRC mouse model resulted in a significant reduction in tumor burden and increased survival. The experimental evidence from mouse colon cancer models and human disease strongly support GPR15 function in immune cell traffic to colonic polyps/tumors to alter the ‘local’ immune contexture to prevent tumor development and/ or mediate tumor resolution.

## Results

### Reduced GPR15 expression and GPR15^+^ immune cells in human colon cancer

To gain insights into GPR15-mediated immune function(s) in colon cancer, we assessed tumor-associated immune environment changes and frequency of GPR15^+^ immune cells in tumor versus surgical tumor margin (STM) of human CRC samples using flow cytometry. STMs were tumor-free as evidenced by H&E as well as clinical pathologist’s assessment and used *in lieu* of healthy controls. (**Fig. 1A**). The core gating strategy for the FACS analysis is outlined in **Supplementary Fig. 1**. We found increased CD45^+^ immune cells (∼2-3 fold) and among CD45^+^ cells, a significant increase only in T cells but not B cells, NK cells, mast cells, or macrophages in the tumor samples compared with STM (**Fig. 1B)**. Moreover, the frequencies of CD4^+^ and CD8^+^ T cells were significantly up-regulated in tumor tissues, with striking increases in IFN*γ*, TNF*α* and IL-17A expression in CD8^+^ T cells and augmented IL-17A expression in CD4^+^ T cells (**Fig. 1C)**. Additional immune features of note included a striking increase in the CD4/CD8 T cell ratio in tumors compared to STM and marked changes in the relative abundance of effector T cell subsets. A comparative analysis revealed a notable shift in the ratio of IFN*γ*^+^ CD8^+^ to IL-17A^+^ CD8^+^ T cell frequencies (Tumor-31.2% vs 1.4% ; STM-17.3% vs 0.4%). The ratios of IFN*γ*^+^ CD8^+^ to TNF*α*^+^ CD8^+^ T cells also showed alteration favoring more TNF*α*^+^ cells while the ratios of TNF*α*^+^ CD8^+^ to IL-17A^+^ CD8^+^ T cells showed no change. A similar trend was observed in the relative frequencies of IFN*γ*^+^ CD4^+^ to IL-17A^+^ CD4^+^ T cells (Tumor 9.7% vs 5.5% ; STM 8.2% vs 1.3%). Interestingly, the ratio of TNF*α*^+^ CD4^+^ to IL-17A^+^ CD4^+^ T cells was also skewed towards more IL-17A^+^ CD4^+^ T cells in tumor vs STM while the ratios of IFN*γ*^+^ CD4^+^ and TNF*α*^+^ CD4^+^ T cells showed a small change towards more TNF*α*. Consistent with previous reports, we also noticed an increase in regulatory CD4^+^ T cells in tumor compared with STM though it did not reach significance. The change in the frequencies of tumor-infiltrating immune subsets observed in our human CRC cohort is reminiscent of tumor immune environments with abundant immunosuppressive effector T cells. To assess if there was a correlation between GPR15 expression and altered immune cell frequencies in our CRC cohort, we examined GPR15 expression in the immune infiltrates from the same human CRC tissues. Our results indicate that GPR15 expression was reduced and frequency of GPR15^+^ CD45^+^ immune cells was decreased in tumor versus STM. Importantly, we noted that GPR15 expression on T cells and frequency of GPR15^+^ CD4^+^ T cells were significantly less in tumors, with a decreasing trend in the frequency of GPR15^+^ CD8^+^ T cells as well (**Fig. 1D)**. GPR15 expression was also significantly decreased in other immune cell subsets, such as natural killer T cells (NKT), macrophages and B cells. Decreased frequencies of GPR15^+^ immune subsets such as NKT, macrophages were observed as well (**Supplementary Fig. 2**). Our data suggest a potential link between differential GPR15 expression and altered composition of both innate and adaptive immune subsets in human CRC depending on disease status (tumor compared to ‘normal’ tumor-margin) with a marked shift towards certain inflammatory, immune-suppressive effector phenotypes.

**Figure 1.**
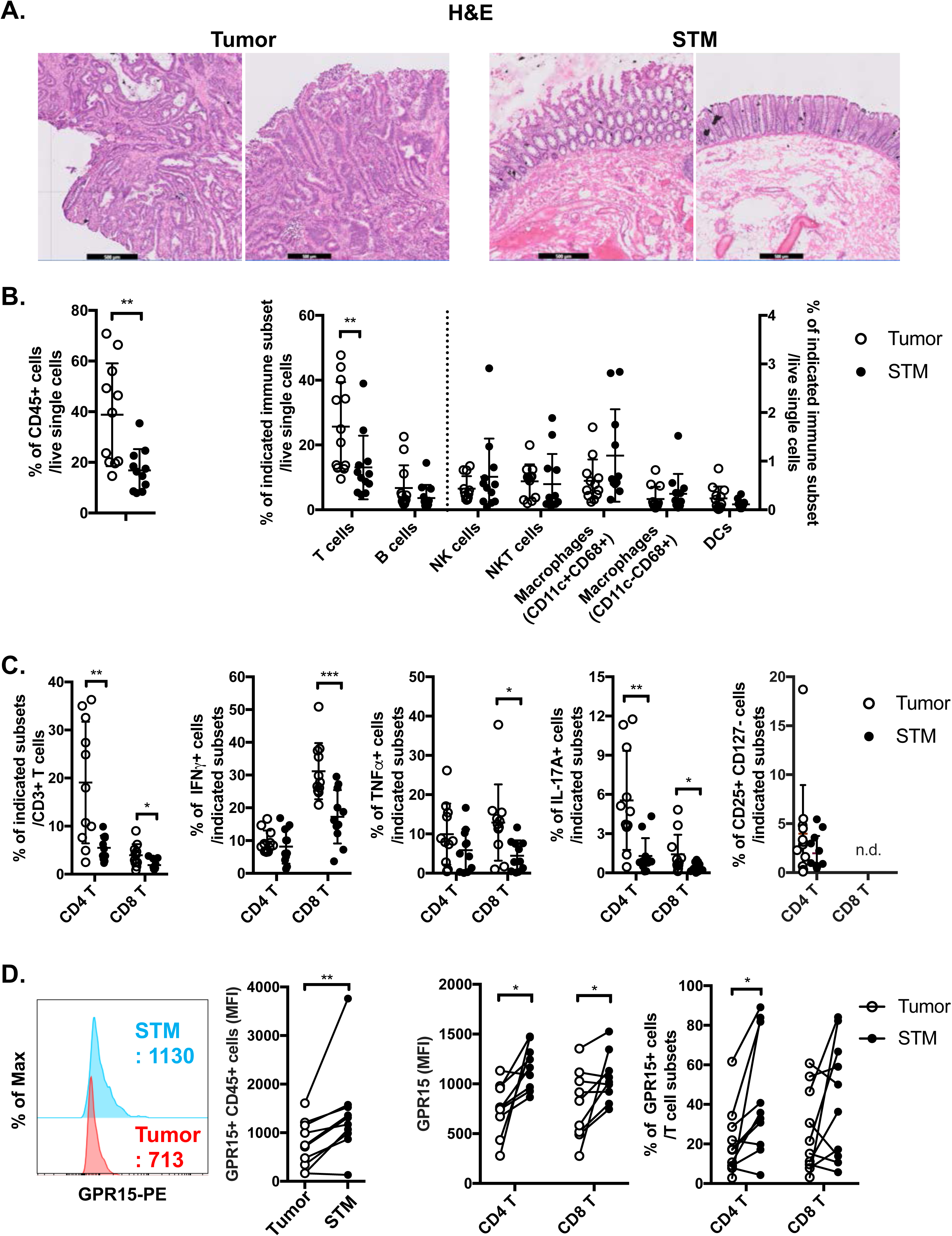
Reduced GPR15 expression and GPR15^+^ immune cells in human colon cancer. **(A)** H&E staining of tissue sections from human CRC, tumor and STM. Scale bar, 500μM. **(B)** Flow cytometry analysis of total CD45^+^ immune cells (left) and innate and adaptive immune cells (right) were compared between tumor and STM tissues. **(C)** Flow cytometry analysis of frequencies of total and cytokine secreting CD4^+^ and CD8^+^ T cells (IFN*γ*, TNF*α*, and IL-17A) in tumor and STM. **(D)** GPR15 expression on immune cells isolated from CRC tissue samples, tumor and STM was analyzed by flow cytometry. Data shown are mean fluorescence intensity (MFI) values for GPR15 and frequency of GPR15^+^ cells in CD45^+^ cells (left), CD4^+^, and CD8^+^ T cells (right). n=10-11. Students *t-test*. \**p*<0.05, \*\**p*<0.01, \*\*\**p*<0.001.

### Lower GPR15 expression is associated with poor survival in human colon cancer

To assess the impact of GPR15 expression and its correlation to patient prognosis in human colon adenocarcinoma (COAD), we analyzed TCGA transcriptomics data using two different publicly available analysis platforms, GEPIA^20^ and TIMER2.0^21^. Our analysis revealed significantly reduced GPR15 mRNA expression in tumors compared to tumor adjacent normal tissue in COAD. Importantly, low GPR15 expression dramatically reduced/shortened patient survival, suggesting that decreased GPR15 expression may be linked to colon cancer disease progression (**Supplementary Fig. 3A, B; top panels**). GPR15L levels were also significantly lower in tumors compared to normal tissue in human COAD though it did not significantly affect patient survival (**Supplementary Fig 3A, B; bottom panels**). Reduced GPR15 expression on tumor-infiltrating T cells in human colon cancer and its impact on disease pathogenesis has not been previously reported and prompted us to investigate its significance using mouse colon cancer models.

### GPR15 deficiency promotes colonic tumors and reduces survival in the AOM-DSS murine model of colitis induced colon cancer

To examine the role of GPR15 in CRC development, we used the well-characterized AOM-DSS induced colitis associated colon cancer (CAC) experimental model to study the course of tumor formation and development in *Gpr15*-sufficient (Het) and *Gpr15*-deficient (knockout; KO) mice^17^ (**Fig. 2A**). The AOM-DSS colon carcinogenesis model recapitulates the multi-step, tumor development sequence and histopathological features of human CRC and is valuable to examine the role of inflammation in colon cancer development. In this model, the colon-tropic carcinogen azoxymethane (AOM) is used in combination with the inflammation-causing agent dextran sodium sulfate (DSS), which erodes the colonic epithelia resulting in severe chronic colitis characterized by bloody diarrhea and tumor formation in the middle and distal colon of murine within 10 weeks ^22–25^. Based on our observation that GPR15^+^ T cells are reduced in the tumor microenvironment in human CRC, we posited that GPR15 deficiency would result in enhanced disease susceptibility and severity in part by influencing T cell traffic to the colon in the AOM-DSS model of CAC. Tumor development was monitored in live mice by endoscopy at week 3, 6 and 9 during the course of AOM-DSS treatment. In line with our hypothesis, we observed severe, widespread inflammation in the colon along with increased incidence of colonic polyps/tumors in the *Gpr15*-KO group compared to *Gpr15*-Het group in both early and late phases of disease development (Day 24-66) post AOM-DSS administration (**Fig 2B and Supplementary Fig 4A**). Importantly, *Gpr15*-KO mice showed significant body weight loss during disease progression (**Fig. 2C**), lower survival rate (**Fig. 2D**), more colonic polyps (of size > 2mm) in the middle and distal colon (measured at week 10; **Fig. 2E, F, Supplementary Fig. 4B, C**), and shorter colon length (measured at week 10; **Fig. 2G**) than *Gpr15*-Het mice. Histology analysis and disease severity scoring after AOM-DSS treatment revealed more severe intestinal inflammation in *Gpr15*-KO compared to *Gpr*15-Het mice, as evidenced by increased necrosis of the epithelial layer, crypt damage, and infiltration of leukocytes in the lamina propria (**Fig. 2H**). *Gpr15*-Het and *Gpr15*-KO mice that were not treated with AOM-DSS showed no polyp formation and had similar survival rates, while C57BL/6 WT mice treated with AOM-DSS demonstrated similar disease course/severity and survival rates as *Gpr15*-Het mice (**Fig. 2C-H** and **Supplementary Fig. 3C**). Our results thus indicate that GPR15 loss led to increased tumor burden, severe disease pathology and poor survival in the AOM-DSS CAC disease model alludes to a protective role of GPR15 in colitis associated colon cancer development.

**Figure 2.**
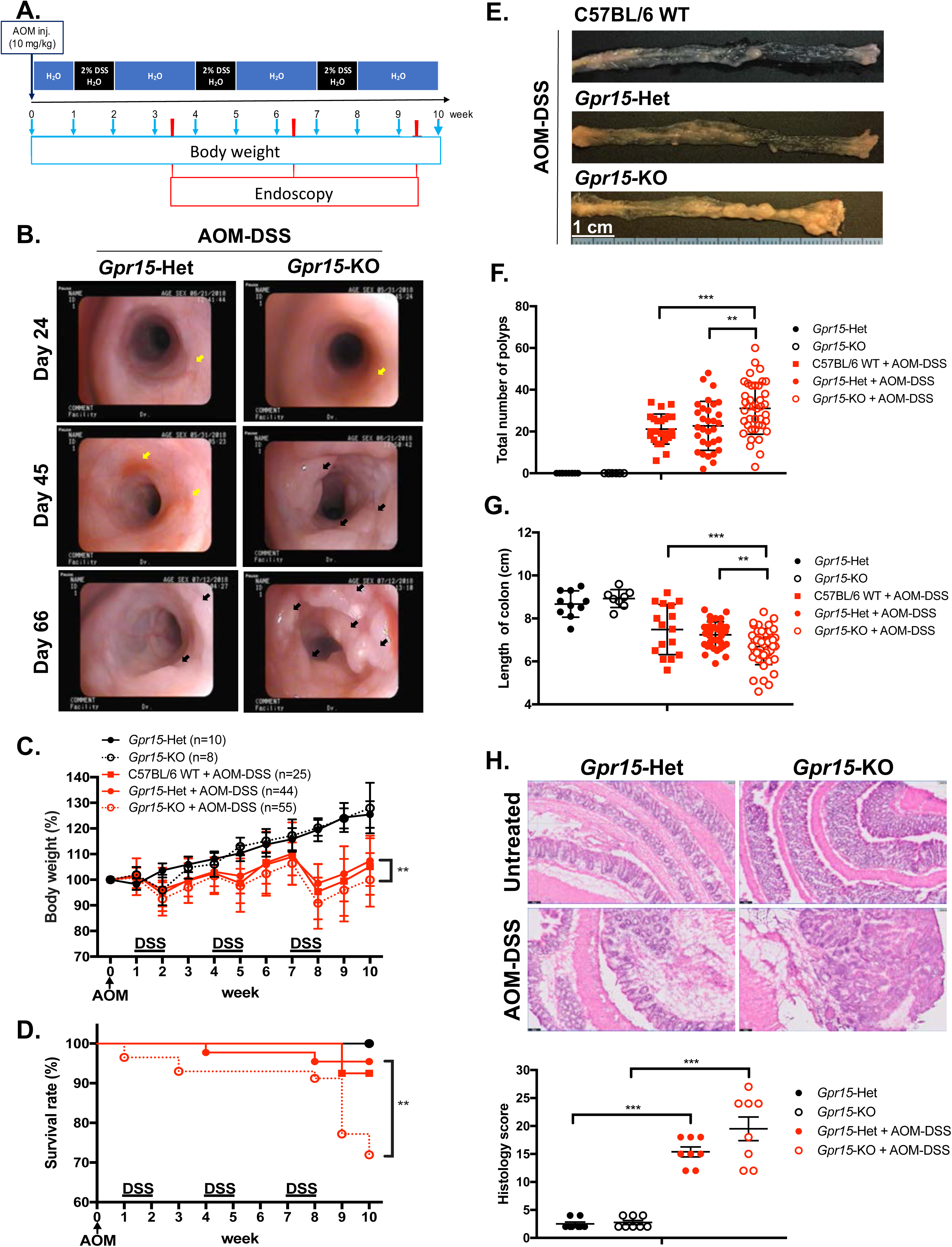
GPR15 deficiency promotes colonic tumors and reduces survival in the AOM-DSS murine model of colitis-induced colon cancer. **(A)** Experimental scheme for the AOM-DSS murine colon cancer model with time points for body weight and endoscopy measurements indicated. **(B)** Representative images of the distal colon in *Gpr15*-Het and KO mice obtained by murine endoscopy on days 24, 45, and 66 during the course of AOM-DSS treatment. Inflammation sites are indicated by yellow arrows, and polyps are indicated by black arrows. **(C)** Bodyweight of untreated and AOM-DSS treated wild type, *Gpr15*-Het and KO mice was measured at different time points as shown in A and reported relative to baseline at start of treatment. Mean ± SD is shown. (**D**) Survival rate, (**E**) representative pictures of the mouse colons, **(F)** the total number of polyps and **(G)** colon length was assessed during the treatment **(C-D)** or after completion of 10-week AOM-DSS treatment **(E-G)** in different mouse groups as indicated. **(C**, **D**, **F**, and **G)**. Data shown is the sum of 5 independent experiments. \*\**p*<0.01, \*\*\**p*<0.001 by Mantel-Cox test or Students *t-test*. **(H)** Representative H&E staining (top) and histological evaluation of disease severity (bottom) of colon sections from *Gpr15-*Het or *Gpr15-*KO mice with or without AOM-DSS treatment (n=8 per group). Scale bar, 100μm. Mean ± SD; by Mann Whitney U-test. \*\*\**p*<0.001.

### GPR15 deficiency alters the immune phenotype and function to favor a pro-tumor environment in the AOM-DSS model of CAC

To delineate GPR15-mediated immune mechanisms involved in tumor suppression in the AOM-DSS CAC disease model, we analyzed the immune cells recovered from the colon in *Gpr15*-Het and *Gpr15*-KO mice at week 10, post AOM-DSS treatment by flow cytometry. Our FACS panel included markers to identify innate and adaptive immune subsets including NK cells, myeloid cells, and T cell subsets (**Supplementary Fig. 5**). We observed a decrease in the total number of T cells and a significant reduction in CD8^+^ T cells in *Gpr15*-KO mice compared to *Gpr15*-Het mice (**Fig. 3A**). To ascertain tumor-specific changes in the immune environment, we performed a comprehensive assessment of immune cell phenotypes and their functional state in large intestine lesions with polyps (LIP) and without polyps (LI) of AOM-DSS treated *Gpr15*-Het and *Gpr15*-KO mice (**Fig. 3B**). We observed that GPR15 deficiency led to significant expansion of regulatory CD8^+^ T cells (CD25^+^ Foxp3^+^) in the LI, and significant increase in frequencies of IL-17A^+^ CD4^+^ and IL-17A^+^ CD8^+^ T cells in LIP of AOM-DSS treated mice. Lower frequencies of IFN*γ*^+^ T cells (both CD4^+^ and CD8^+^) were noted in LI or LIP of AOM-DSS mice regardless of GPR15 expression (**Fig. 3C**). Furthermore, we noted a marked increase in the frequency of myeloid-derived suppressor cells (MDSC; CD11b^+^ Gr-1^+^) in the LI and LIP (**Fig. 3D**). Interestingly, the colonic polyps (LIP) in the AOM-DSS treated *Gpr15*-KO mice showed distinct (e.g., increased MDSCs) as well as shared (e.g., increased IL-17A^+^ CD4^+^ and IL-17A^+^ CD8^+^ T cells) immune features of human CRC with reduced GPR15 expression. The decreased infiltration of CD8^+^ T cells and expanded MDSC population observed in the tumors (LIP) of AOM-DSS treated *Gpr15*-KO mice are common denominators in tumor formation and progression in several cancers including CRC^26^. Our findings in the AOM-DSS CAC model lend credence to GPR15 involvement in recruitment of T cells with tumor-suppressive function to the colon and aiding in the evolution of an immune environment unfavorable for tumor growth.

**Figure 3.**
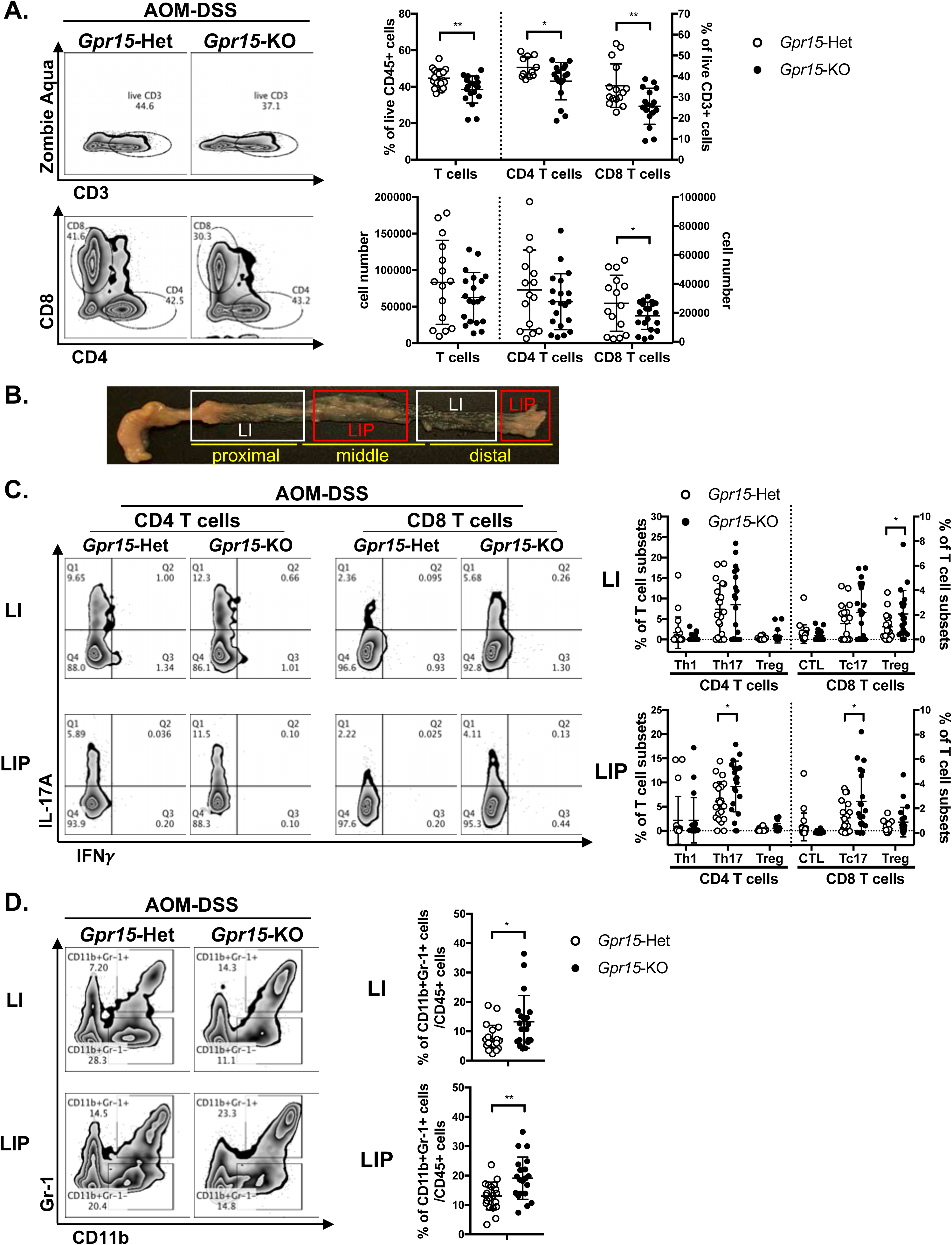
GPR15 deficiency alters immune cell phenotypes and effector T cell populations in the colon tumor microenvironment in the AOM-DSS model. **(A)** Representative flow cytometry plots of CD3^+^ T cells and their subpopulations, CD4^+^ and CD8^+^ T cells (left) isolated from the colon of AOM-DSS treated *Gpr15*-Het and KO mice as indicated. Frequencies (Top, right) and absolute cell numbers (bottom right) of CD3^+^, CD4^+,^ and CD8^+^ T cells are shown. **(B)** Representative picture showing the large intestine without polyps (LI) and the large intestine with polyps (LIP). **(C)** Representative flow cytometry plots of IL-17A or IFN*γ* expressing CD4^+^ and CD8^+^ T cells from LI and LIP (left) and frequencies of T cell subsets (Th1, Th17, and Treg) in LI and LIP (right). **(D)** Representative plots (left) and frequencies (right) of MDSCs from LI and LIP of AOM-DSS administered *Gpr15-*Het and *Gpr15-*KO mice. **(A**, **C**, and **D)** Frequencies are shown as dot plots, mean ± SD. Students *t-test* \**p*<0.05, \*\**p*<0.01.

Our further investigation of the immune cell frequencies in the small intestine and secondary lymphoid organs, such as spleen (SP), Peyer’s patches (PP), mesenteric lymph nodes (MLN), and peripheral (axillary, brachial, inguinal, and sciatic) lymph nodes (PLN) revealed the presence of diverse immune populations at varying frequencies with some notable differences between the tissues of AOM-DSS treated *Gpr15*-Het and *Gpr15*-KO mice (**Supplementary Fig. 6**). Activated dendritic cells (MHC-II^+^ F4/80^-^ CD11c^+^), CD4^+^ and CD8^+^ T cells were significantly decreased while macrophages (F4/80^+^ CD11c^-^) were expanded in the MLN of AOM-DSS treated *Gpr15*-KO mice. Myeloid-derived suppressor cells (MDSCs, CD11b^+^ Gr-1^+^) were significantly increased while CD8^+^ T cells were decreased in the spleen. Furthermore, the absence of GPR15 altered the frequencies of diverse effector subsets in the different tissues as measured by cytokine expression. Thus, the expression of pro-inflammatory cytokines, such as TNF*α*, IFN*γ* and IL-12, was decreased on CD4^+^ T cells, CD8^+^ T cells and DCs (F4/80^-^ CD11c^+^) present in the lymphoid organs (SP and PLN) of AOM-DSS treated *Gpr15*-KO mice (**Supplementary Fig. 7**). Moreover, anti-inflammation cytokine, TGF*β*, was elevated on CD4^+^ Foxp3^+^ T cells in the MLN of AOM-DSS administrated *Gpr15*-KO mice. The above-mentioned differences in immune infiltrates were specifically observed in the context of GPR15 deficiency in AOM-DSS treated mice and was not observed in untreated, *Gpr15*-KO mice (**Supplementary Fig. 7**). Our immune analysis in the secondary lymphoid organs thus reveal systemic, tumor-favoring immune signatures in *Gpr15*-KO mice (summarized in Supplementary **Table S2**) that reflect the changes in the ‘local’ colonic tumor immune environment (eg. increased MDSC, CD4^+^ subsets) and further highlight an important role for GPR15-mediated immune mechanisms in colon cancer development.

### GPR15 agonism elicits anti-tumor effects in the murine MC38 colon cancer model

Our findings in human CRC and murine AOM-DSS colon carcinogenesis model highlight a novel function of GPR15 as a tumor-suppressor by modulating the immune microenvironment in colon cancer. To further delineate the role and explore the translational relevance of our findings, we examined if GPR15 agonism by GPR15L administration would elicit an anti-tumor effect by influencing immune cell phenotypes and their functionality in the MC38-murine CRC model. The MC38 cell-line was originally derived from a grade-III colon adenocarcinoma that was chemically induced in a female C57BL/6 mouse^27^ and is a well-established CRC model for investigating anti-cancer immunity and immunotherapy^28–30^. MC38 cells were injected subcutaneously in *Gpr15*-Het and - KO mice, and tumor development was also monitored following concomitant intra-tumoral injection of GPR15L or vehicle when tumors reached 4-5 mm in size (Day 4 post-transplant). Consistent with the results in our AOM-DSS model, the tumors in the *Gpr15*-KO mice were significantly bigger than in *Gpr15*-Het mice (PBS treated group; **Fig. 4A, B**). Importantly, GPR15L administration resulted in dramatic shrinkage in tumor size in *Gpr15*-Het but not in *Gpr15*-KO mice, affirming a specific function of the GPR15-GPR15L axis in mitigating colon tumor growth. Flow cytometry analysis of GPR15-GPR15L signaling dependent changes in immune cell populations in the tumors revealed significant expansion of CD45^+^ T cells, CD4^+^ and CD8^+^ T cells, in particular TNF*α*^+^ T cell subsets, in *Gpr15*-Het but not in *Gpr15*-KO mice treated with GPR15L (**Supplementary Fig. 8 and Fig. 4C, D**). The MC38 tumors had low frequencies of IFN*γ*^+^ T cell infiltrates, which remain unchanged in Het and KO, similar to our observations in the colonic polyps of AOM-DSS treated mice (PBS group; **Fig 4D**). Our results indicate that GPR15-GPR15L axis impairs tumor growth by facilitating the infiltration of ‘cytolytic’ T cells into the tumor microenvironment in the MC38 CRC model. This is supported by the finding that TNF*α*^+^ CD4^+^ and CD8^+^ T cells were reduced in *Gpr15*-KO compared to *Gpr15*-Het mice with PBS treatment. Furthermore, we noticed increased frequency of TNF*α*^+^ NK cells, IL-12^+^, and IFN*γ*^+^ NKT cells, TNF*α*^+^ DCs, and macrophages in the GPR15L-injected tumors isolated from *Gpr15*-Het mice (**Supplementary Fig. 9**). Whether these changes are a consequence of direct or indirect GPR15 function warrants further investigation. No significant changes were observed in IL-12^+^ NK cells, IL-17A^+^ NK and NKT cells, IL-1*β*^+^, IL-6^+^, IL-10^+^, and TGF*β*^+^ dendritic cells (F4/80^-^ CD11c^+^) and CD206^+^ macrophages. In sum, our studies provide ‘proof-of-concept’ for the therapeutic targeting of GPR15-GPR15L axis in CRC treatment; intra-tumoral administration of GPR15L attenuates tumor growth in a GPR15-dependent manner by shaping an anti-tumor immune environment enriched in cytolytic T, NK and NKT cells.

**Figure 4.**
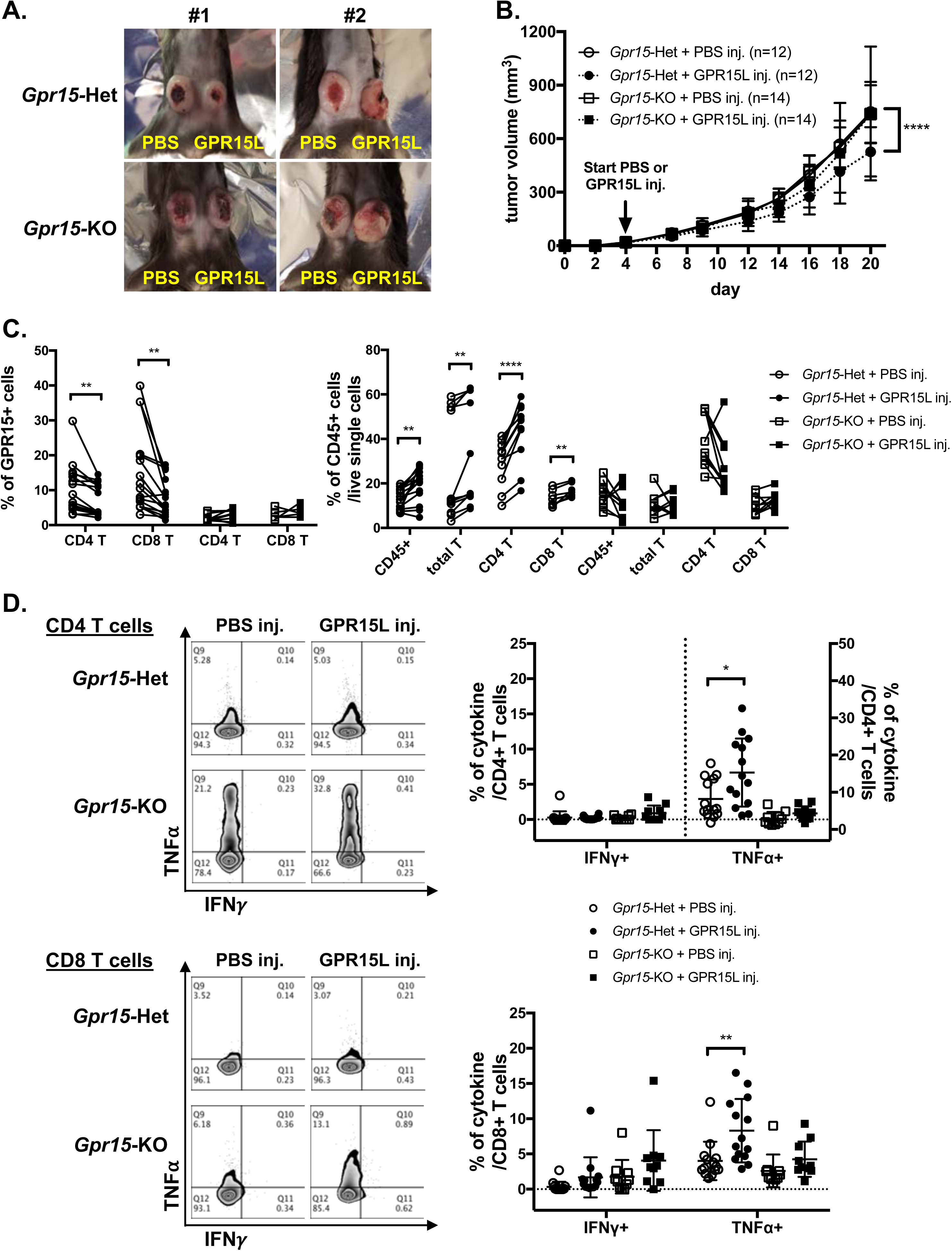
GPR15L treatment facilitates anti-tumor effects in the murine MC38 colon cancer model. **(A)** Representative pictures of PBS and GPR15L-treated tumors (day 21) in *Gpr15*-Het and *Gpr15*-KO mice. **(B)** Tumor growth rates in *Gpr15*-Het and *Gpr15*-KO mice with PBS or GPR15L treatment as indicated. Data shown is mean ± SD. Two-way ANOVA \*\*\*\**p*<0.0001. **(C)** Flow cytometry analysis of GPR15 expression on CD4^+^ and CD8^+^ T cells (left) and frequency of CD45^+^ immune cells and T cell subsets (right) in tumors isolated from different treatment groups as indicated. **(D)** Representative flow cytometry plots (left) and frequencies (right) of IFN*γ* and TNF*α* expressing CD4^+^ T or CD8^+^ T cells in tumors of different treatment groups. Data shown in **C-D** is mean ± SD. Students *t-test* \**p*<0.05, \*\**p*<0.01, \*\*\*\**p*<0.0001.

## Discussion

We previously identified an important function of GPR15 in T cell traffic to the inflamed colon and colitis pathogenesis. In the present study, we describe a first and novel function of GPR15 in colon cancer: GPR15 plays a crucial role in the colonic recruitment of immune cells and shaping of the colon immune microenvironment to mediate a protective, tumor-mitigating effect. This is demonstrated by our *in vivo* studies in the AOM-DSS murine model of CAC, where GPR15 deficiency resulted in a significant reduction of T cells with tumor-suppressive function in the colon, such as effector CD8^+^ T cells and the ability of GPR15L administration to shrink established MC38 tumors in *GPR15*-sufficient but not in *GPR15*-deficient mice. In support of a tumor-protective function of GPR15 in human CRC, GPR15 expression as well as GPR15^+^ immune cells were significantly reduced in human colon tumors and lower GPR15 expression correlated with decreased survival of COAD patients. Our studies offer a strong rationale for modulation of GPR15-GPR15L axis as a colon-specific approach to treat CRC by enhancing effector immune cell responses culminating in tumor rejection.

Our analysis of immune infiltrates in human colon cancer and AOM-DSS mouse CAC model indicated that GPR15 expression may modulate the relative abundance of innate and adaptive immune cell subsets as well as cytokine secreting, effector T cell subsets in the colonic tumor microenvironment (TME) in favor of tumor regression. In our human CRC samples, T cells were the predominant population in tumor tissues compared to STM among immune infiltrates analyzed and among T cells, a higher frequency of CD4^+^ T cells compared to CD8^+^ T cells was noted (19.1% vs 3.9%). The ‘T cell rich’ TME in our patient cohort shows consensus with previous reports as well as studies where higher ratios of CD4/CD8 T cells have been reported in human colon cancer samples^31, 32^. Emerging evidence suggests a link between increased T cell infiltration especially of CD3^+^ T cells and favorable patient prognosis. However, the balance between pathogenic versus protective T cells and their interactions in the TME is a critical factor governing clinical outcomes^8, 33^. Reduced GPR15 expression and frequency of GPR15^+^ immune cells in the TME correlated with an enrichment of certain inflammatory and immunosuppressive phenotypes in our patient cohort. This was evidenced by significantly increased frequencies of IL-17A secreting CD4^+^ and CD8^+^ T cells, and a trend towards increased CD25^+^ CD127^-^ regulatory CD4^+^ T cells in the tumors compared to uninvolved colonic tissue (STM). Furthermore, assessment of frequencies and relative levels of other effector T cell populations in tumor versus STM showed a notable shift in the ratio of IFN*γ*^+^ to IL-17A^+^ T cell frequencies (on both CD4^+^ and CD8^+^ T cell subsets). Interestingly, the ratio of TNF*α*^+^ CD4^+^ and IL-17A^+^ CD4^+^ was also skewed towards more IL17A^+^ CD4^+^ in tumor compared with STM. Higher proportion of Th17 cells have been reported in tumor as well as blood of CRC patients and is associated with tumor progression and a poor prognosis^34^. Interestingly, Amicarella *et al*^35^ demonstrated using human CRC samples that tumor infiltration of IL-17^+^ T cells *per se* was not predictive of patient survival. Rather, the spatial location of IL17^+^ T cells within tumors influenced prognosis; intra-epithelial but not stromal Th17 cells was positively correlated with improved survival. Whether GPR15 has a role in spatial distribution of effector cells within tumor regions to establish unique, intra-tumoral immune niches is currently not known and warrants future investigation.

In the AOM-DSS CAC model, GPR15 deficiency was similarly associated with unique alterations in the composition of immune subsets in the tumor (LIP) and non-tumor (LI) regions of the colon as well as in secondary lymphoid organs. We noted a marked overall decrease in tumor infiltrating CD8^+^ T cells and expansion of myeloid (CD11b^+^ Gr1-1^+^) populations. Of significance was a striking increase in the frequency of IL-17^+^ CD4^+^ (Th17) cells and IL-17^+^ CD8^+^ T cells (Tc17) in the LIP but not in LI of *Gpr15-*KO mice. Interestingly, the colonic polyps (LIP) in the AOM-DSS treated *Gpr15-*KO mice showed distinct (eg. expansion of MDSCs) as well as shared (eg. increased IL-17^+^ CD4^+^ and IL-17^+^ CD8^+^ T cells) immune features of human CRC with reduced GPR15 expression. In contrast to our human CRC data, the frequency of IFN*γ*^+^ CD4^+^ Th1 cells, IFN*γ*^+^ CD8^+^ T cells (CTL) and TNF*α*^+^ T cells (data not shown) were low and unaltered in the LI and LIP of AOM-DSS treated *Gpr15-*KO compared with *Gpr15-*Het mice. The patient cohort we analyzed was not stratified based on factors such as tumor mutations, IBD history, which could differentially affect the nature and composition of tumor infiltrating immune cells. It is thus possible that the observed differences in immune cell composition could be due to heterogeneity and molecular changes unique to the human CRC samples compared to AOM-DSS mouse model and underscore the potential of the TME to influence the infiltration of diverse GPR15^+^ immune cells.

Our results in the MC38 murine colon cancer model demonstrate the potential of GPR15L treatment to alleviate tumor burden in a GPR15-dependent manner. The tumor-suppressing capability of the GPR15-GPR15L axis was associated with a significant influx of CD4^+^ and CD8^+^ T cells into the tumor, in particular TNF*α* secreting T cells. Increased frequencies of TNF*α*^+^ NK cells, TNF*α*^+^ NKT cells and IFN*γ*^+^ NKT cells were also observed following GPR15L treatment of tumors in *Gpr15*-Het. Anti-tumor effects of NK and Type1 NKT in CRC have been reported^36^ and likely contribute to direct or indirect GPR15-GPR15L dependent negative regulation of tumor growth. Unlike the observations in the AOM-DSS model, no alterations in IFN*γ*^+^ IL-17A^+^ T cell frequencies were observed across treatment groups. Thus, there might be differences in GPR15-dependent infiltration of effector CD4^+^ and CD8^+^ T cells depending on the tumor location and subsequent changes in the TME (subcutaneous vs colon). Furthermore, it is possible that GPR15L effects on colonic tumor growth may involve both GPR15-dependent and independent mechanisms and warrants further studies. In this respect, it is worth noting that GPR15L has recently been reported to interact with SUSD2 to inhibit colon cancer cell growth via G1 arrest, which is induced by down-regulating cyclin D1 and cyclin-dependent kinase 6 (CDK6)^37^. Intriguingly, we noted reduced GPR15 expression on tumor infiltrating CD4^+^ and CD8^+^ T cells in GPR15L-treated *Gpr15*-Het mice compared to the PBS-treated *Gpr15*-Het mice. This could be due to downregulation of surface GPR15 upon GPR15L binding. Indeed, a recent report shows that GPR15L facilitates GPR15 endocytosis using GPR15-expressing cell lines^38^. Additional studies are required to understand GPR15L-dependent regulation of surface GPR15 levels on immune cells and its role in GPR15-mediated immune mechanisms in health and inflammatory diseases such as colitis and colon cancer to realize the therapeutic potential of GPR15-GPR15L axis in these indications.

Our current findings on GPR15 expression and function in colon cancer are supported/strengthened by a recent pan-cancer study of GPR15 expression and mutational pattern using data from TCGA and GTex databases that revealed GPR15 was hypermutated and its expression significantly downregulated in colon adenocarcinoma^39^. Importantly, there was a positive correlation between GPR15 expression, CD4^+^ T cell infiltration and patient survival. In the same study, integrated network analysis on GPR15 and GO enrichment analysis revealed potential intersections with proteins involved in mitogenic signaling, cell-cycle regulation and links to immune function in different cancers including colon cancer. The study thus provides a glimpse of possible effects that can be mediated by GPR15 in colon cancer beyond its classical function as a T-cell trafficking protein. Another study on mRNA and miRNA signatures in human COAD patient samples also reported reduced GPR15 expression in tumors and potential miRNA mediated regulation of GPR15 levels in COAD^40^.

The immune environment in CRC is highly heterogenous and complex, evolves during disease progression and poses significant challenges for development of effective immunotherapies. Prominent among the cellular immune mediators in the anti-tumor response are cytotoxic lymphocytes, in particular (primarily) effector CD8^+^ T cells. They are supported in this endeavor by NK cells, Th1-like type I NKT cells and CD4^+^ IFN*γ*^+^ Th1 cells. There is a growing appreciation for defects in trafficking of immune cells (eg. homing deficit of anti-tumor effector T cells) to tumor sites as a key impediment to the success of adoptive T cell transfer-based therapies in cancers including CRC^41^. Our current findings that the T cell homing receptor, GPR15 can augment the infiltration of tumor-fighting effector T cells and alter the TME in colon cancer lesions in pre-clinical models hold therapeutic promise in immunotherapy-based tumor eradication using GPR15 agonists and improving patient survival.

## Materials and Methods

### Patient samples

Colon resection samples were obtained from patients undergoing adenocarcinoma indicated colectomies with approval by the Stanford University Institutional Review Board and after obtaining informed patient consent. The clinicopathological features of the patient samples are summarized in Table S1. After removal of the serosa, mesenteric fat and muscularis externa, leukocytes were isolated from the lamina propria as described^42^. Cells were resuspended in RPMI containing 5% FBS and re-stimulated with 50 ng/ml phorbol 12-myristate 13-acetate (PMA, Sigma) plus 1 μg/ml ionomycin (Sigma) in the presence of 5 μg/ml Brefeldin A (BioLegend) for 4 hours at 37℃ prior to staining for flow cytometry analysis.

### Analysis of human COAD TCGA datasets

GPR15 and GPR15L expression in human COAD TCGA cohorts was analyzed using TIMER2.0 (http://timer.cistrome.org) and GEPIA (http://gepia.cancer-pku.cn) webservers^20, 21^. Briefly, TIMER2.0 ‘Gene_DE’ module was used to assess differential expression of GPR15 and GPR15L in tumor and adjacent normal tissues in TCGA cancer types collected from GDAC firehose website. TIMER2.0 extracts raw counts and Transcripts Per Million (TPM) from RSEM results. Additional analysis of GPR15 and GPR15L expression in human COAD tumor tissue and normal tissue as well as its association with patient survival was done using GEPIA webserver which utilizes RNAseq data from TCGA and GTex projects (source: UCSC Xena project; http://xena.ucsc.edu).

### In vivo studies

C57BL/6 and B6.SJL mouse strains were purchased from Jackson Laboratory and bred in-house. GPR15-GFP mice, generated as described^17^, were backcrossed 10 times onto the C57BL/6 backgrounds before use in experiments. *Gpr15*-Het (*Gpr15^gfp/+^*) and *Gpr15*-KO (*Gpr15^gfp/gfp^*) mice used in our studies are ‘knock-in’ mice in which endogenous GPR15 was replaced with the sequence for GFP. Animals were maintained in accordance with the US National Institutes of Health guidelines, and experiments were approved by Stanford University Institutional Animal Care and Use Committee.

To induce colonic tumors using AOM-DSS, 8 weeks old mice were injected intraperitoneally with the procarcinogen azoxymethane (AOM) (10 mg/kg of body weight, Sigma-Aldrich). After 1 week, the mice received drinking water supplemented with 2% DSS (MP Biomedicals; MW, 36-50 kDa) for 7 days, followed by 2 weeks of regular water. The DSS administration was repeated twice with 2% DSS in drinking water^43–45^. Body weight was measured once a week for 10 weeks relative to baseline during AOM-DSS treatment. Serial endoscopy was performed at day 24, 45 and 66 in live mice to visualize the distal region of the colon and monitor tumor growth. Mice were euthanized 10 weeks after the AOM administration, and the colon length and the number of polyps were recorded. Immune cells were isolated from the mice colons and assessed by flow cytometry.

The murine colon adenocarcinoma cell line, MC38, was kindly provided by Jeanette Baker, Stanford University, USA and maintained in DMEM supplemented with 10% FBS, 1mM L-Glutamine, Penicillin (100 µg/ml), Streptomycin (100μg/ml) and 2-Mercaptoethanol (50 μM), at 37 °C in a 5% CO_2_ incubator. For tumor induction, MC38 cells (5 × 10^5^ cells/site) were implanted subcutaneously (s.c.) in the left and right flank of female *GPR15*-Het or *GPR15*-KO mice. When the tumor diameter reached 4-5 mm (on day 4), mice were administrated intratumorally (i.t.) with 50 μl of PBS (left flank) or murine recombinant GPR15L procured from PeproTech (right flank, 2.5 μg in PBS/injection) 3 times per week for 2 weeks. Tumor size was measured 3 times per week using a digital caliper and mice were euthanized when tumors reached a size of 1.7 cm in each dimension. Immune cells were isolated from tumor tissues using the cell isolation methods described above and stained for flow cytometry analysis.

### Mouse colonoscopy

Severity and size of tumors in distal part of the colon were determined by micro-colonoscopy endoscopy at day 24, day 45 and day 66 of AOM-DSS treatment as described^46^. Mice were anesthetized using 2% isoflurane (Forane, Baxter, Deerfield, Il). The colorectal lumen of the sedated animals was cleaned with 37°C warm phosphate-buffered saline (PBS) and pellets removed. The endoscope (G110074, Pentax) was then inserted slowly into the anus of the animals. The video processing, light, and airflow were delivered by the video processor (EPK-1000, Pentax). The white-light endoscopy was recorded using GrabBee software on a laptop connected using a VGA to HDMI connector.

### Cell isolation

Single cell suspension of leukocytes was prepared from mouse spleen, Peyer’s patches, mesenteric and peripheral (including inguinal, brachial sciatic and axillary) lymph nodes by mechanically dispersing the tissues through a stainless steel 100 μm wire mesh into Hank’s buffered salt solution (HBSS; Mediatech, Inc.) containing 2% BCS. Splenic red blood cells were lysed with Red Blood Cell Lysing buffer (Sigma-Aldrich). Intraepithelial and lamina propria lymphocytes were isolated from the small intestine and colon as described previously^47^. After excision of the Peyer’s patches, the intestines were cut into ∼0.5 cm pieces and cleansed in HEPES-buffered RPMI medium containing 5% BCS. To isolate lamina propria (LP) lymphocytes, the tissue pieces were incubated in 2mM EDTA for 15 min (x 2 rounds) followed by 3 rounds of collagenase (0.7 mg/ml) digestion at 37° C (20 min each), and the supernatant was collected, pooled, and washed. Centrifugation in a 40% and 70% Percoll step-gradient produced enriched colon lamina propria lymphocytes at the interface which were collected, washed, and stained for flow cytometry analysis.

### Flow cytometry and antibodies

For intracellular cytokine assays, cells were re-stimulated with PMA (50 ng/ml) and ionomycin (1 μg/ml) in the presence of Brefeldin A (5 μg/ml) for 4 hours at 37℃ prior to flow cytometry staining. Dead cells were first excluded using Zombie Aqua™ Fixable Viability Kit (BioLegend). Cells were then incubated with fluorescence conjugated antibodies in HBSS containing 2% BCS at 4℃ for 30 minutes to stain surface markers. Surface-stained cells were permeabilized with Cytofix/Cytoperm kit (BD Biosciences) and incubated with fluorescence labeled antibodies to different cytokines in HBSS containing 2% BCS at 4° C for 30 minutes. The following antibodies were used for flow cytometry in the mouse studies: CCR1 (S15040E), CCR9 (CW-1.2), CD3 (17A2, 145-2C11), CD4 (RM4-5, GK1.5), CD11b (M1/70), CD11c (N418), CD19 (1D3/CD19), CD25 (PC61), CD45 (30-F11), CD103 (2E7), CD117 (ACK2), CD206 (C068C2), F4/80 (BM8), Gr-1 (RB6-8C5), IL-4 (11B11), IL10 (JES5-16E2), IL-22 (Poly5164), MHC-II(I-A/I-E) (M5/114.15.2), NK1.1 (PK135), TGF-b (TW7-16B4), TNF-a (MP6-XT22) from BioLegend; CD8 (53-6.7), CD45 (30-F11), IFN-g (XMG1.2), IL10 (JES5-16E2), IL-12(p40/p70) (C15.6), IL-17A (TC11-18H10) from BD Biosciences; Foxp3 (IC8214A) from R&D Systems. For staining human cells from colon tissues, the following antibodies were used: CCR1 (5F10B29), CD3 (UCHT1), CD8 (SK1, RPA-T8), CD11c (3.9), CD19 (HIB19), CD25 (BC96), CD45RO (UCHL1), CD68 (Y1/82A), CD117 (104D2), CD127 (A019D5), CD206 (15-2), GPR15 (SA302A10), IL-10 (JES3-19F1), IL-17A (BL168), IFN-g (B27, 4S.B3), TNF-a (Mab11), Perforin (B-D48), mouse IgG2a isotype control (MOPC-173) from BioLegend; CD4 (RPA-T4), CD206 (19.2) from BD Biosciences; GPR15 (FAB3654P) from R&D Systems. Data was acquired on an LSRII or Fortessa cytometer (BD Biosciences) and analyzed with FlowJo software (FlowJo, LLC). Cytokine secretion was calculated by subtracting the signal from unstimulated cells from that of the stimulated cells.

### Histology of colon tissues

Haematoxylin and Eosin (H&E) staining was performed by Histology Service Center (HIC) at Stanford University. Histological scoring was performed in a blinded fashion by two independent observers under the guidance of histology scoring method as described elsewhere^45^. Histology score = (Grade of Inflammation + Grade of extent + Grade of crypt damage) x Grade of percent involvement.

### Statistical analysis

Student’s *t-test*, Mann–Whitney U test, Mantel-Cox test or ANOVA was performed using Prism (GraphPad) to analyze all experimental data unless otherwise stated. \**p*<0.05; **, *p*<0.01; \*\*\**p*< 0.001.

## Acknowledgments

We thank patients and donors who provided precious human colon tissues for this study. We thank Yujun Yang, Allison Ji, and Yi Wei for their technical assistance, such as mouse genotyping and animal care. We also thank all of the members of Habtezion Laboratory for their helpful comments and suggestions. The study was in part supported by the Ann and Bill Swindells Charitable Trust, and NIH R01 grant, DK101119 to A.H.

## Author contribution

H.N. designed the study, performed all experiments, analyzed and interpreted data, and wrote the manuscript; B.L. contributed to mouse experiments, data analysis, and edited the manuscript; G.S. contributed to data analysis and wrote the manuscript; S.K. contributed to mouse experiments and histology scoring; S.R. performed mouse colonoscopy; D.M. provided technical assistance for *in vivo* studies. A.H. designed and supervised the studies, provided guidance, analyzed data and wrote the manuscript. All the authors had the opportunity to discuss the results, review, and comment on the final manuscript.

## Supplementary Material

**Supplementary Figure 1.**
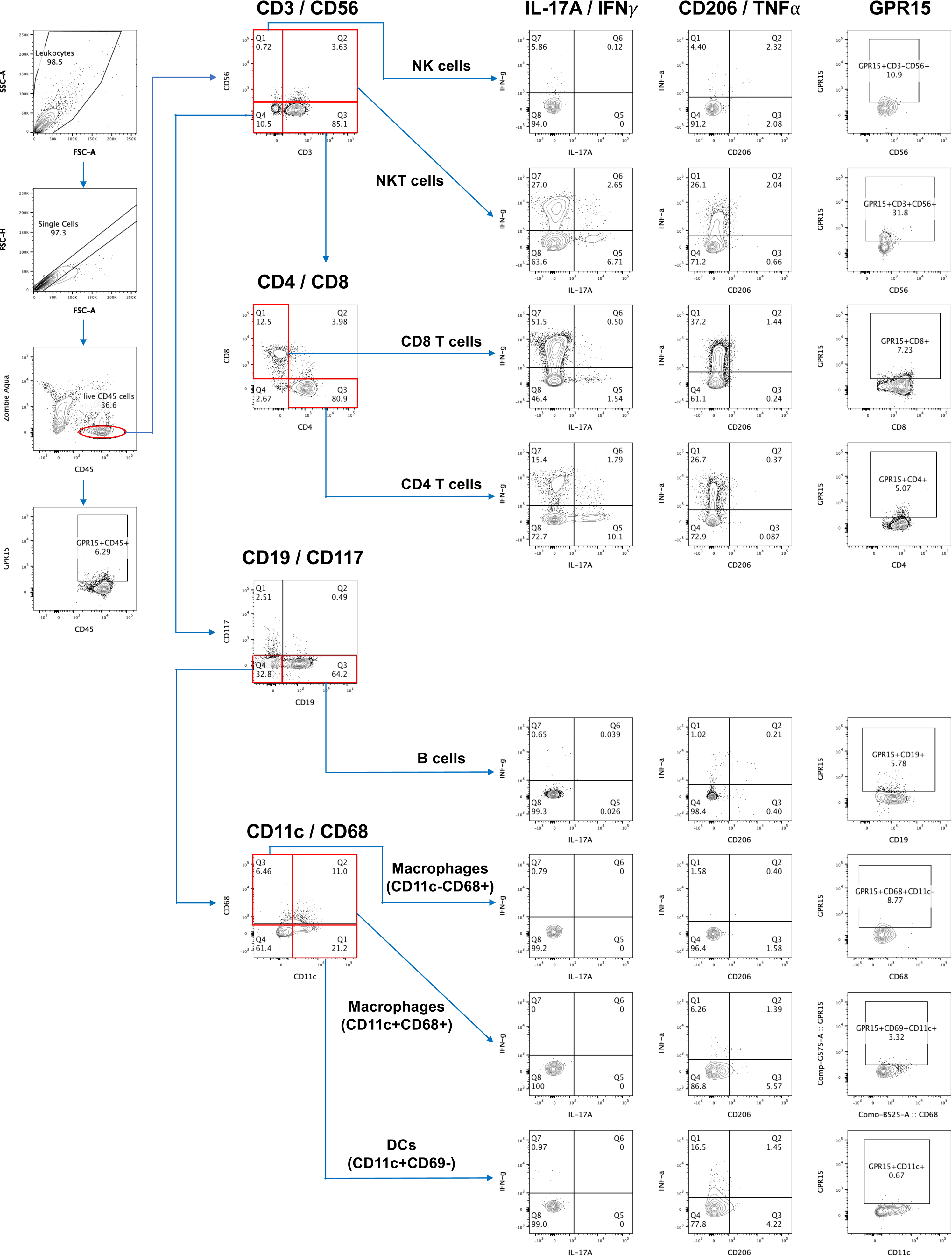
Core gating strategy for analysis of immune cell subsets and functional phenotypes in human CRC patient samples.

**Supplementary Figure 2.**
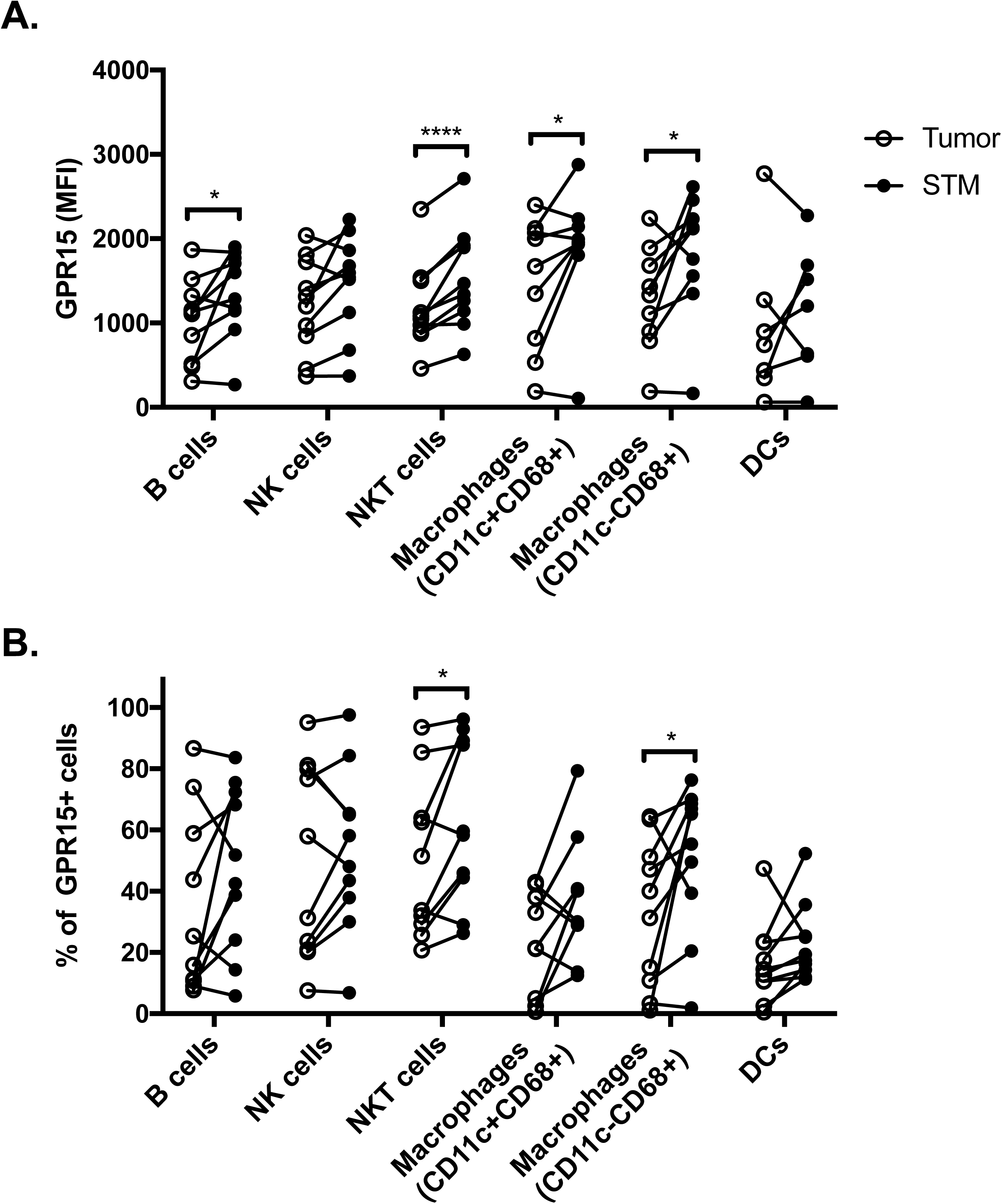
Differential GPR15 expression in innate and adaptive immune cell subsets in human CRC. **(A)** GPR15 levels (MFI) on immune cells and (**B**) frequency of GPR15^+^ immune cells in tumor versus STM of human CRC samples were assessed by flow cytometry. n=10. Students *t-test*; \**p*<0.05, \*\*\*\**p*<0.0001.

**Supplementary Figure 3.**
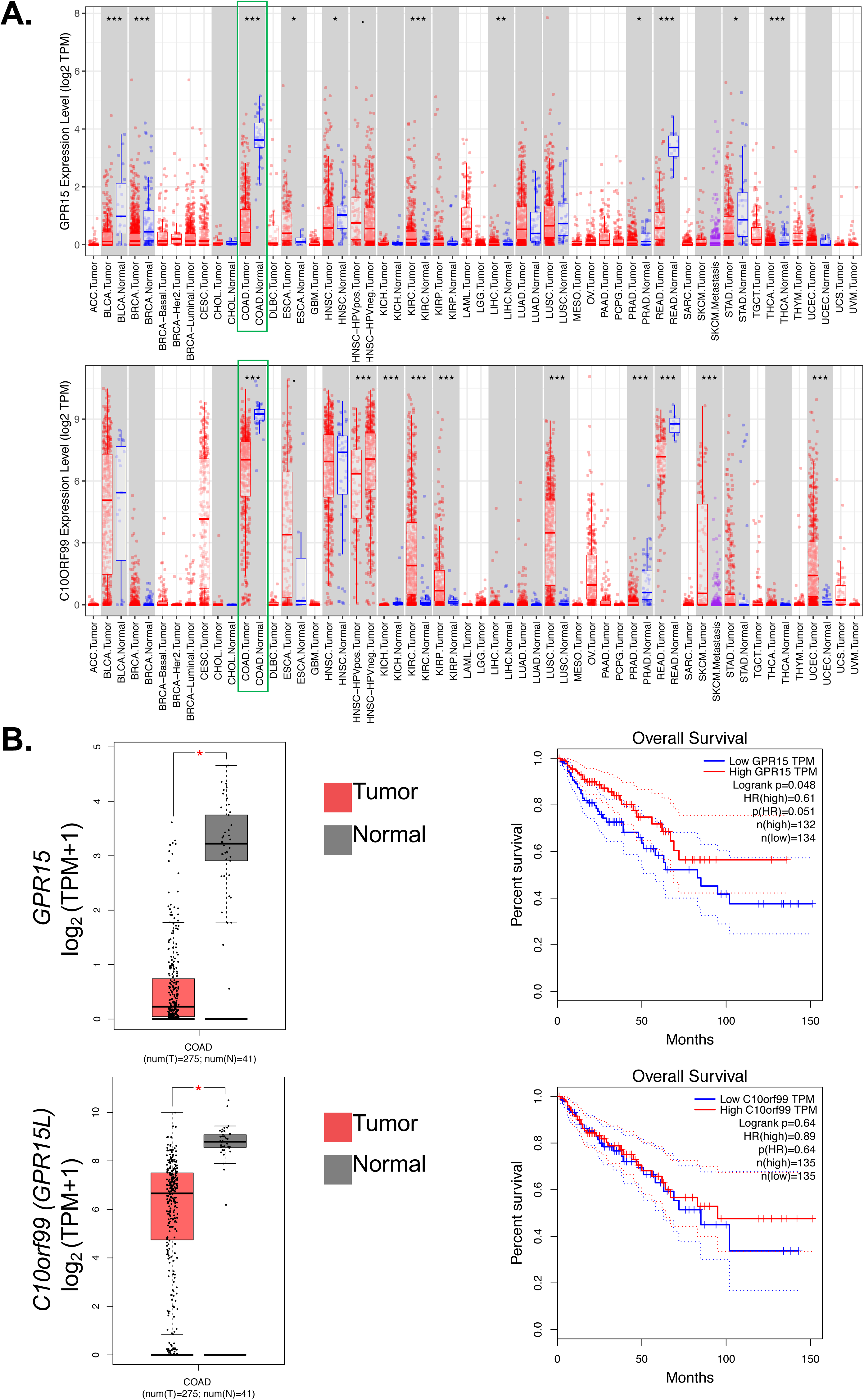
Expression of *GPR15* and *GPR15L* is reduced in human colon adenocarcinoma (COAD) and associated with poor patient prognosis. **(A)** Analysis of *GPR15* and *GPR15L (C10orf99)* expression in tumor versus normal tissue using TCGA datasets and TIMER2.0 analysis platform in different human cancers. Expression in COAD (n=) is highlighted in green **(B)** Analysis of *GPR15* and *GPR15L* expression (left panels) and its correlation to patient survival (right panels). Tumor and paired normal tissue in TCGA-COAD datasets were analyzed with GEPIA analysis platform. \**p*<0.05, \*\**p*<0.01, \*\*\**p*<0.001.

**Supplementary Figure 4.**
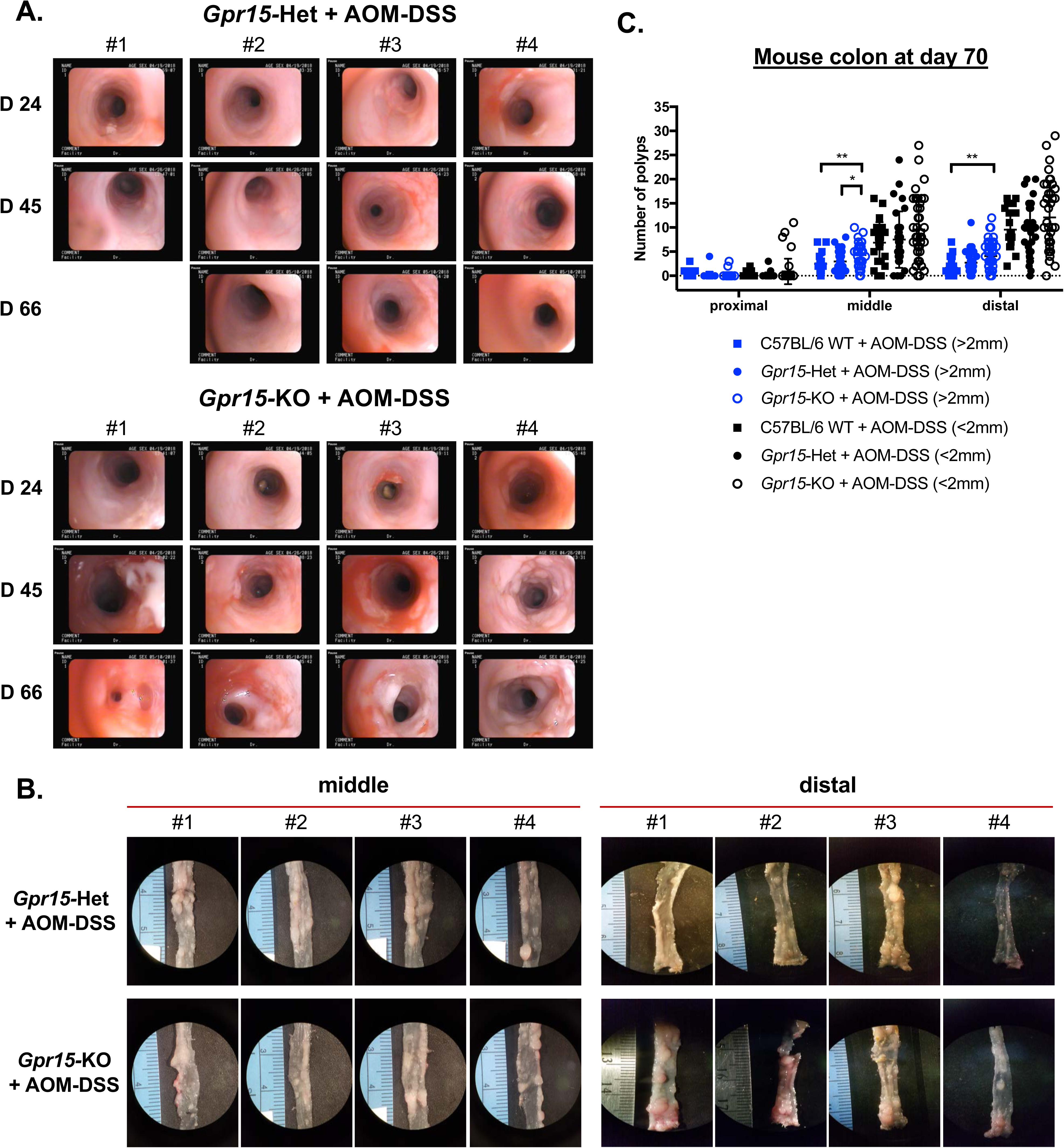
*Gpr15* deficiency accelerates colitis-associated colon cancer development. **(A)** Endoscopic images of the distal colon on day 24, day 45 and day 66 of AOM-DSS treated *Gpr15-*Het and *Gpr15-*KO mice. **(B)** Representative images of polyps/tumors in the middle and distal colon of *Gpr15*-Het and *GPR15*-KO at week 10, post AOM-DSS treatment. **(C)** Quantitation of polyp size and numbers in the proximal, middle and distal colon of WT, *Gpr15*-Het and *Gpr15*-KO mice at week 10 post AOM-DSS administration. Blue; polyp size>2mm, Black; polyp size<2mm. n=24-31. Students *t-test*. \**p*<0.05, \*\**p*<0.01.

**Supplementary Figure 5.**
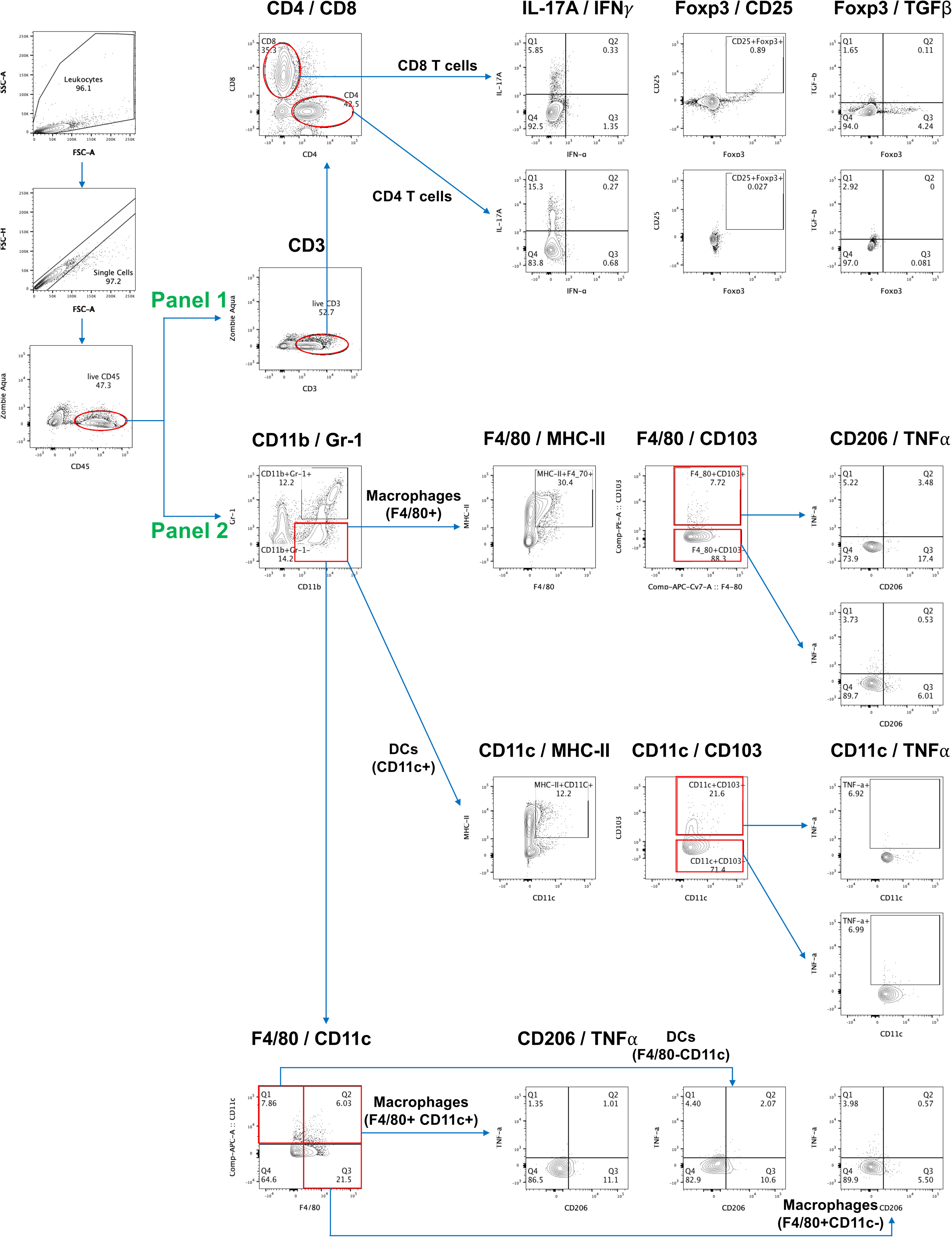
Core gating strategy for analysis of immune cell subsets and functional phenotypes in the large intestine and secondary lymphoid organs in the AOM-DSS mouse CAC model.

**Supplementary Figure 6.**
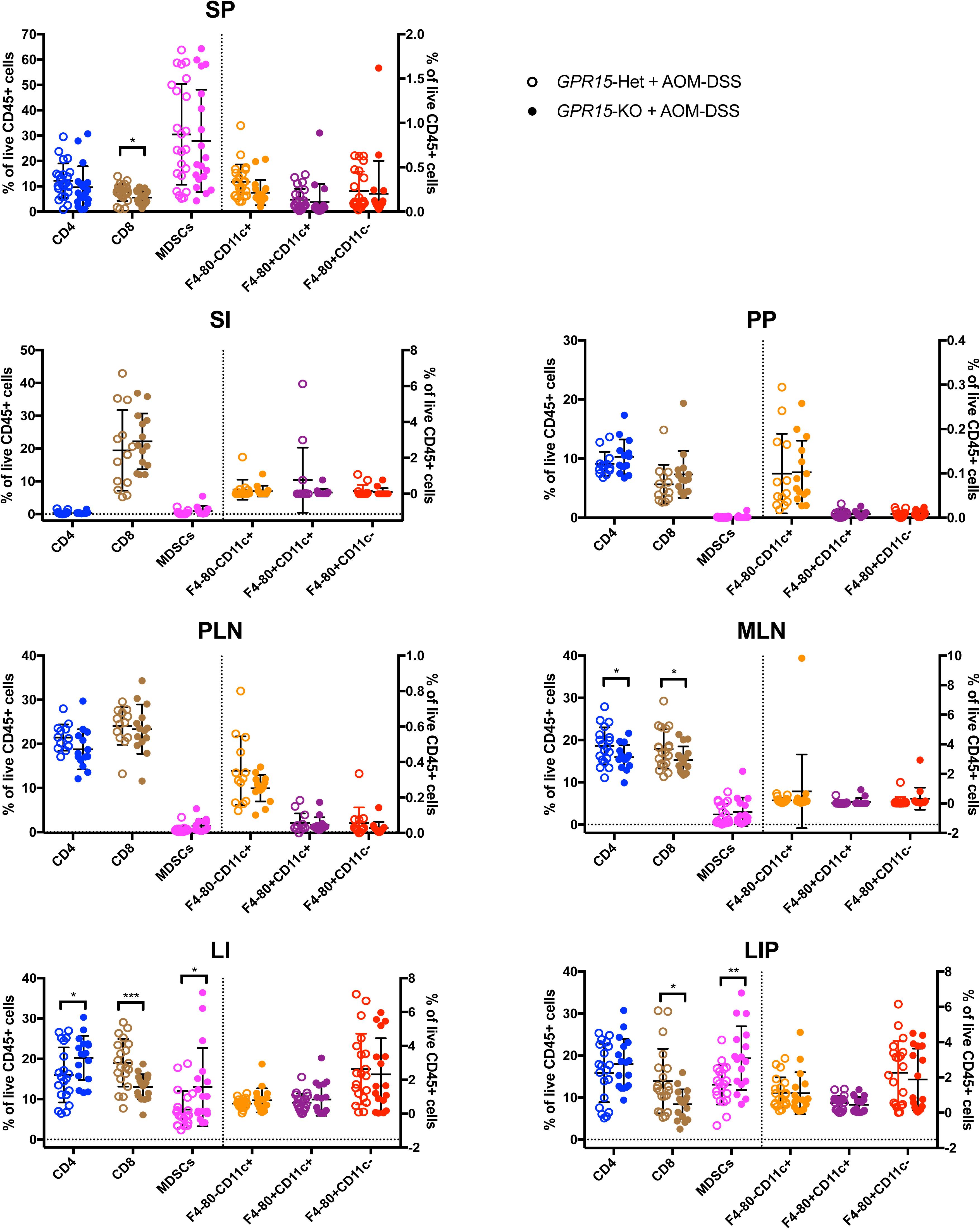
Alteration of immune subset frequencies in the large intestine and secondary lymphoid organs in AOM-DSS treated *Gpr15*-Het and *Gpr15*-KO mice. Frequencies of immune subsets, as indicated, analyzed in spleen (SP), small intestine (SI), peyer’s patches (PP), peripheral lymph nodes (PLN), mesenteric lymph nodes (MLN), large intestine (LI), and large intestine with polyps (LIP). n=19-23. Students *t-test*. \**p*<0.05, \*\**p*<0.01, \*\*\**p*<0.001.

**Supplementary Figure 7.**
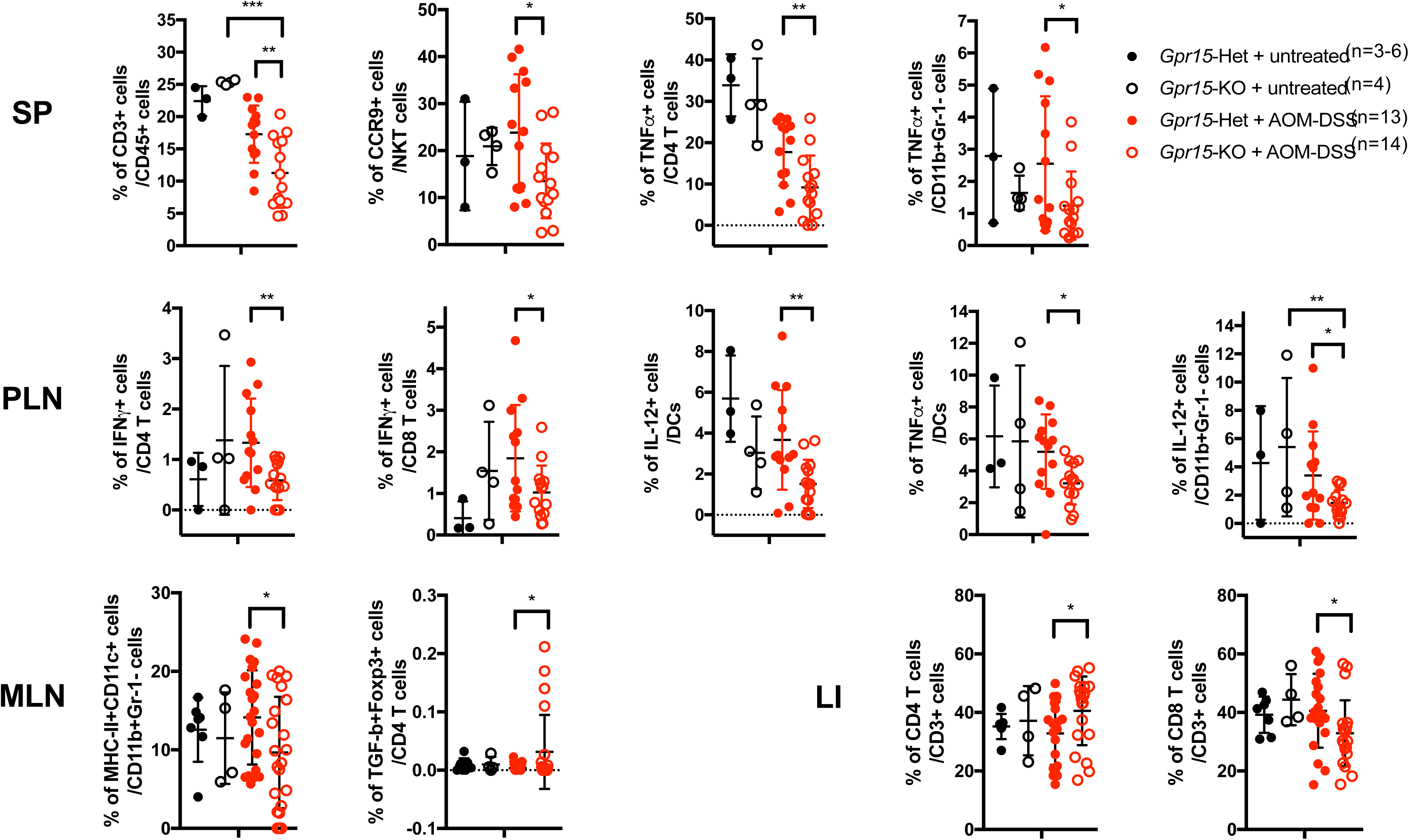
Alteration of T cell and myeloid cell subsets frequencies in the large intestine and secondary lymphoid organs in AOM-DSS treated *Gpr15*-Het and KO mice. Frequencies of immune subsets, as indicated, analyzed in spleen (SP), peripheral lymph nodes (PLN), mesenteric lymph nodes (MLN), and large intestine (LI). Mean ± SD; n=13-14. Students *t-test*. \**p*<0.05, \*\**p*<0.01, \*\*\**p*<0.001.

**Supplementary Figure 8.**
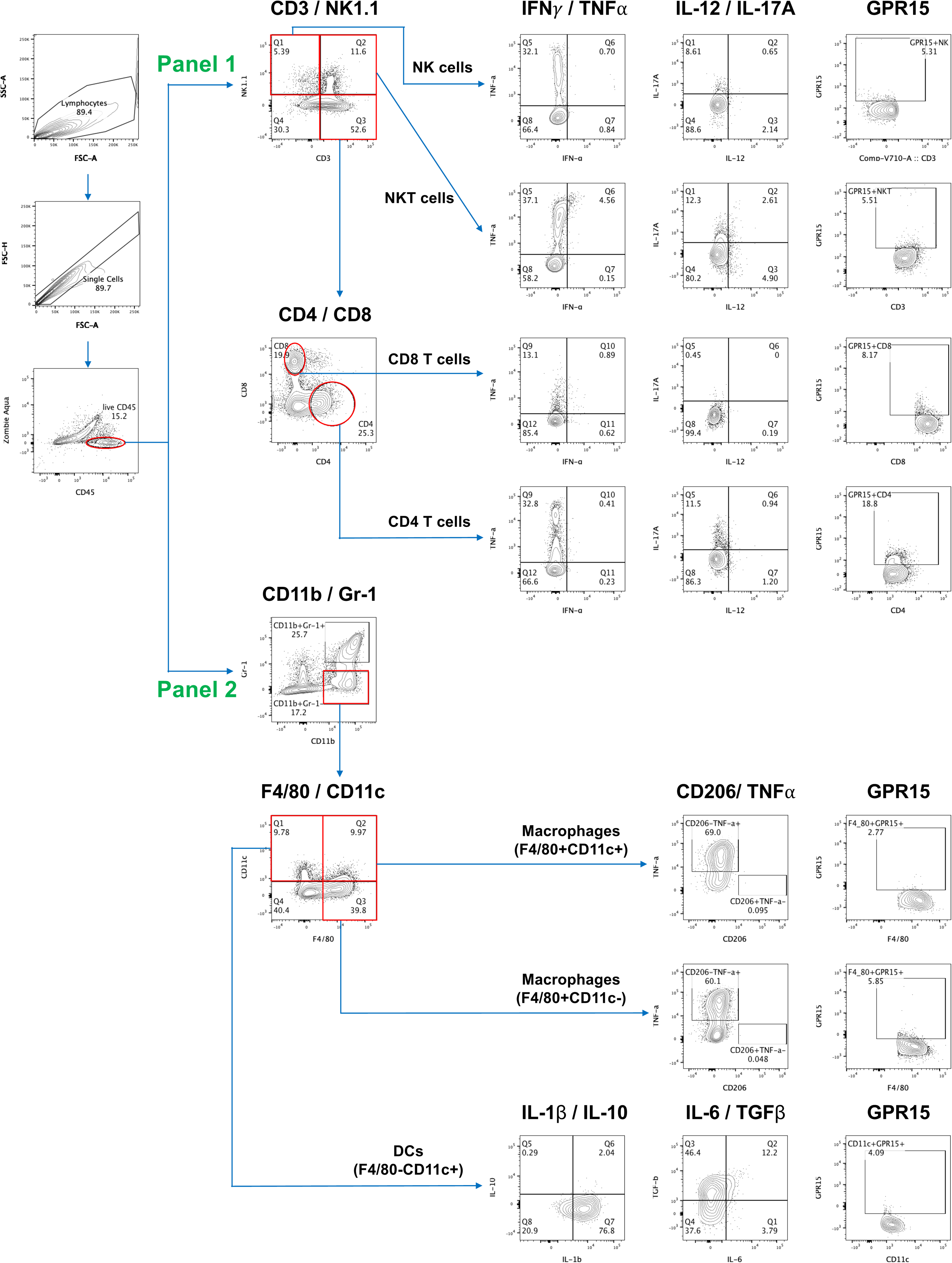
Core gating strategy for analysis of immune cell subsets and functional phenotypes in the large intestine and secondary lymphoid organs in the MC38 mouse CRC model.

**Supplemental Figure 9.**
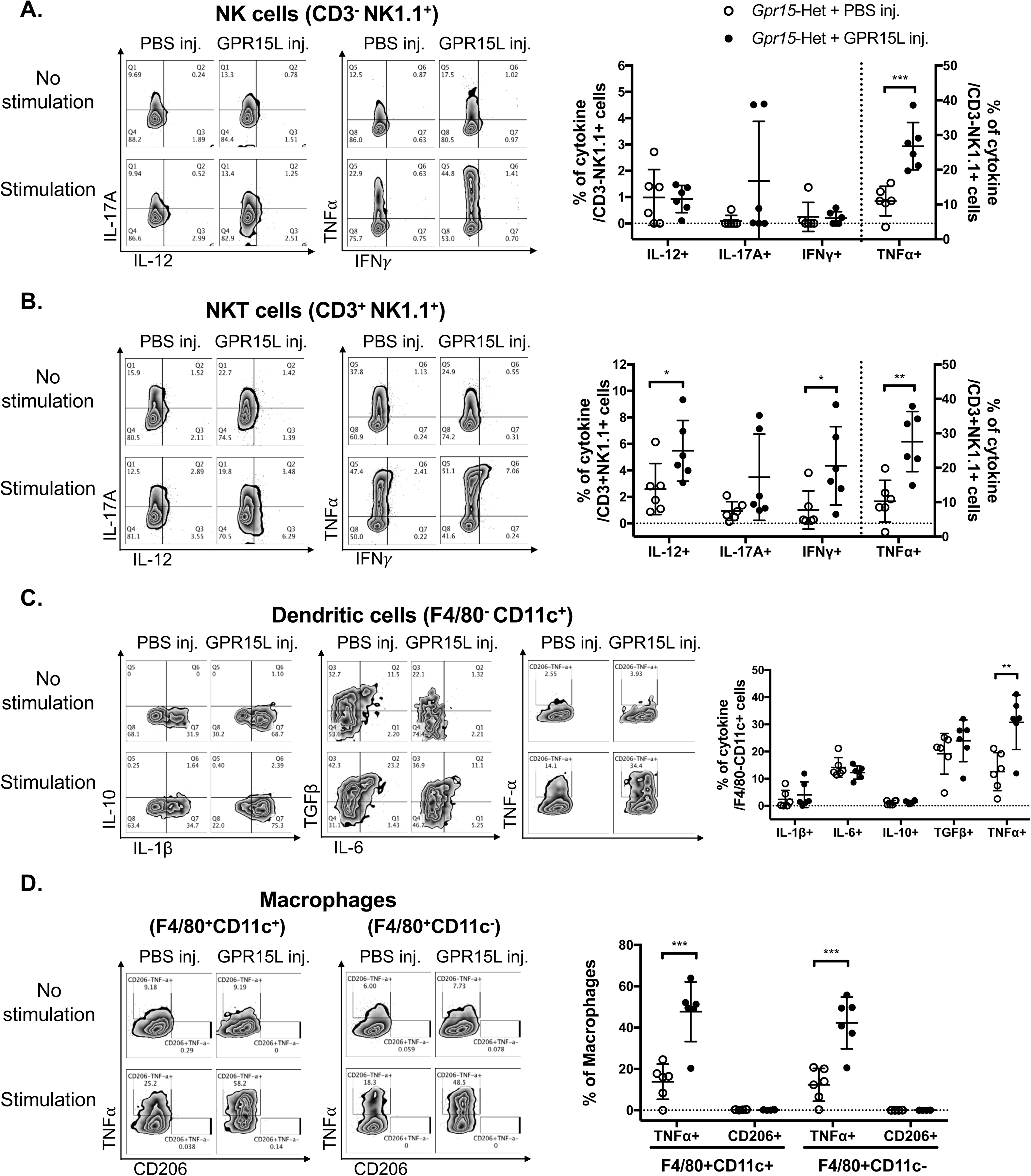
Frequencies of effector subsets in PBS and GPR15L injected MC38 tumors in *Gpr15*-Het and *Gpr15-KO* mice. Representative FACS plots (left) and quantitation (right) of cytokine expression in NK cells (A), NKT cells (B), dendritic cells (C), and macrophages (D). Mean ± SD; n=6. Students t-test. \**p*<0.05, \*\**p*<0.01, \*\*\**p*<0.001.

**Table S1.**
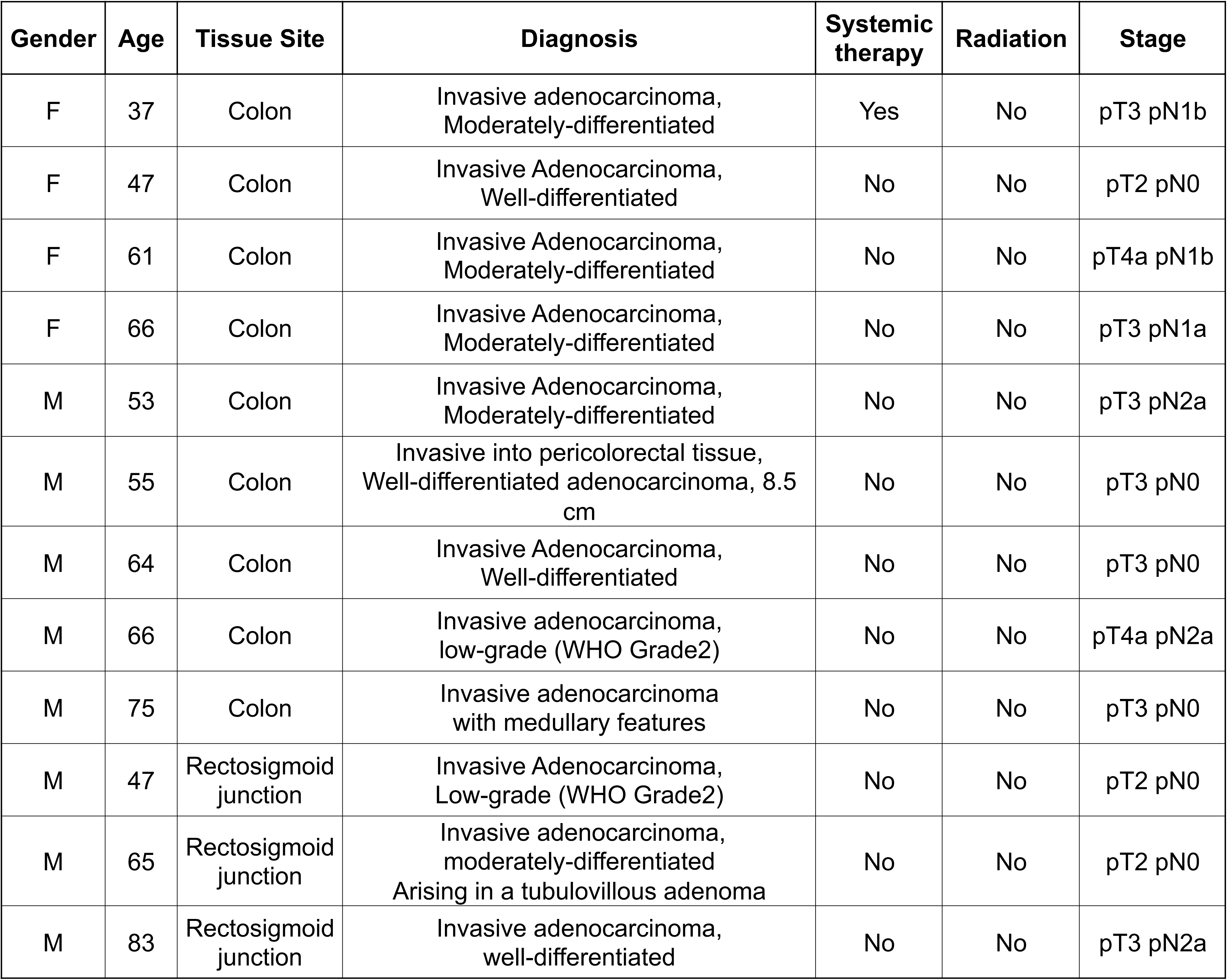
Clinicopathological characteristics of human CRC patients used in the study.

**Table S2.**
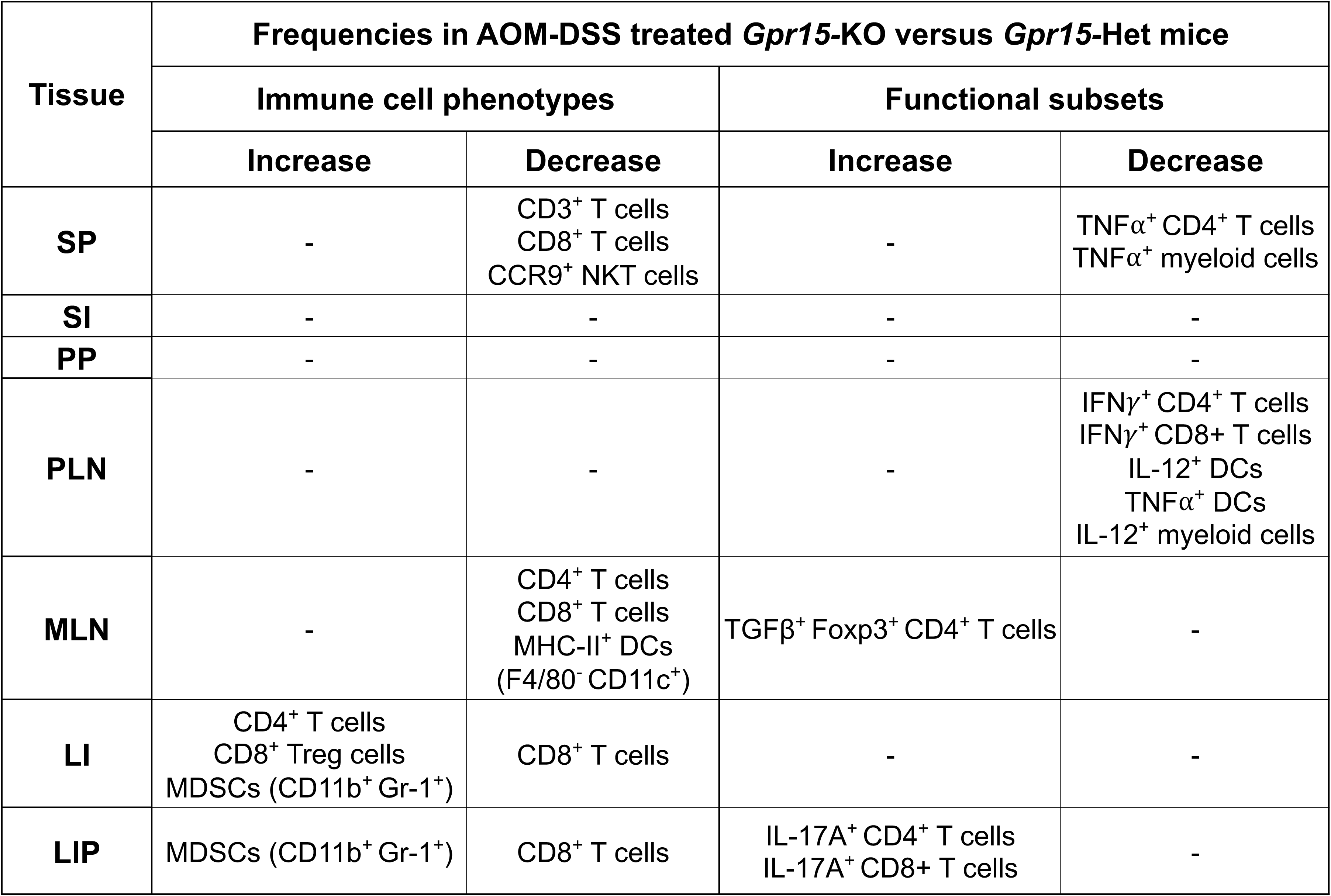
Summary of changes in frequencies of T cells and myeloid cells in different tissues in AOM-DSS treated *Gpr15*-KO versus *Gpr15*-KO mice.

| Tissue | Frequencies in AOM-DSS treated *Gpr15*-KO versus *Gpr15*-Het mice |  |  |  |
| --- | --- | --- | --- | --- |
|  | Immune cell phenotypes |  | Functional subsets |  |
|  | Increase | Decrease | Increase | Decrease |
| SP | - | CD3<sup>+</sup> T cells<br>CD8<sup>+</sup> T cells<br>CCR9<sup>+</sup> NKT cells | - | TNFα<sup>+</sup> CD4<sup>+</sup> T cells<br>TNFα<sup>+</sup> myeloid cells |
| SI | - | - | - | - |
| PP | - | - | - | - |
| PLN | - | - | - | IFNγ<sup>+</sup> CD4<sup>+</sup> T cells<br>IFNγ<sup>+</sup> CD8<sup>+</sup> T cells<br>IL-12<sup>+</sup> DCs<br>TNFα<sup>+</sup> DCs<br>IL-12<sup>+</sup> myeloid cells |
| MLN | - | CD4<sup>+</sup> T cells<br>CD8<sup>+</sup> T cells<br>MHC-II<sup>+</sup> DCs<br>(F4/80<sup>-</sup> CD11c<sup>+</sup>) | TGFβ<sup>+</sup> Foxp3<sup>+</sup> CD4<sup>+</sup> T cells | - |
| LI | CD4<sup>+</sup> T cells<br>CD8<sup>+</sup> Treg cells<br>MDSCs (CD11b<sup>+</sup> Gr-1<sup>+</sup>) | CD8<sup>+</sup> T cells | - | - |
| LIP | MDSCs (CD11b<sup>+</sup> Gr-1<sup>+</sup>) | CD8<sup>+</sup> T cells | IL-17A<sup>+</sup> CD4<sup>+</sup> T cells<br>IL-17A<sup>+</sup> CD8<sup>+</sup> T cells | - |

## Notes

### Competing Interest Statement

The authors have declared no competing interest.

